# Stromal fibroblast activation and inflammation in frozen shoulder

**DOI:** 10.1101/453894

**Authors:** Moeed Akbar, Michael McLean, Emma Garcia-Melchor, Lindsay AN Crowe, Paul McMillan, Umberto G Fazzi, David Martin, Angus Arthur, James H Reilly, Iain B McInnes, Neal L Millar

**Author notes:** These authors contributed equally (Moeed Akbar, Michael McLean). Correspondence: Neal L Millar PhD FRCSEd(Tr&Ortho), Senior Clinical Lecturer in Orthopaedics, Honorary Consultant Orthopaedic Surgeon, Institute of Infection, Immunity and Inflammation, College of Medicine, Veterinary and Life Sciences, University of Glasgow, 120 University Avenue, Glasgow G12 8TA.

## Abstract

**Introduction:** Frozen shoulder is a common, fibro-proliferative disease characterised by the insidious onset of pain and progressively restricted range of shoulder movement. Despite the prevalence of this disease, there is limited understanding of the molecular mechanisms underpinning the pathogenesis of this debilitating disease. Previous studies have identified increased myofibroblast differentiation and proliferation, immune cell influx and dysregulated cytokine production. We hypothesised that subpopulations within the stromal compartment may take on an activated phenotype, thus initiating the inflammatory processes observed in frozen shoulder. Therefore, we sought to evaluate the presence and possible pathogenic role of known stromal activation proteins in Frozen shoulder,

**Methods:** Shoulder capsule samples were collected from 10 patients with idiopathic frozen shoulder and 10 patients undergoing shoulder stabilisation surgery. Stromal activation marker expression (CD248, CD146, VCAM and PDPN, FAP) was quantified using immunohistochemistry. Control and diseased fibroblasts were cultured for in vitro studies from capsule biopsies from instability and frozen shoulder surgeries, respectively. The inflammatory profile and effects of IL-1β upon diseased and control fibroblasts was assessed using ELISA, immunohistochemistry and qPCR.

**Results:** Immunohistochemistry demonstrated increased expression of stromal activation markers CD248, CD146, VCAM and PDPN in the frozen shoulder group compared with control (p < 0.05). Fibroblasts cultured from diseased capsule produced elevated levels of inflammatory protein (IL-6, IL-8 & CCL-20) in comparison to control fibroblasts. Exposing control fibroblasts to an inflammatory stimuli, (IL-1ß) significantly increased stromal activation marker transcript and protein expression (CD248, *PDPN and VCAM).*

**Conclusions:** These results show that stromal fibroblasts have an activated phenotype in frozen shoulder and this is associated with inflammatory cytokine dysregulation. Furthermore, it supports the hypothesis that activated stromal fibroblasts may be involved in regulating the inflammatory and fibrotic processes involved in this disease.

## Introduction

The fibroproliferative disorder, frozen shoulder is a multifactorial disease in which patients present with limited active and passive shoulder movement^35^. Despite a prevalence of 2-5% and known risk factors including diabetes, thyroid disease, stroke and autoimmune diseases there remains a significant lack of understanding of the molecular mechanisms underpinning this common fibrotic disorder which affects many recreational sporting individuals^31^.

Recent studies have demonstrated altered expression of immune cells, inflammatory mediators and fibrosis associated cytokines in frozen shoulder. Chronic inflammatory cells including mast cells, T and B cells and macrophages have been identified in shoulder capsule biopsies from patients with frozen shoulder^18^ while diseased capsule showed dysregulated cytokines such as IL-1β, IL-6 and TNF-α, which are known to drive inflammatory/matrix interactions^27^ including fibroblast activation and dysregulated collagen synthesis. Additionally, we have shown the cytokines interleukin 33 (IL-33) as an alarmin in early tendinopathy^24^ whereby when released from the resident tenocyte has the ability to modulate inflammatory/matrix crosstalk and thus is likely to be important in the balance between reparation and degeneration in tissue repair. Subsequently we discovered increased levels of the alarmins IL33 and high-mobility group protein B1 (HMGB1)^7^, which are known to not only initiate but amplify and sustain the inflammatory processes in frozen shoulder. This reinforces the concept of immune involvement in triggering, regulating and remission of inflammation in frozen shoulder^16^.

Pathological shoulder capsule tissue has demonstrated myofibroblast differentiation and proliferation in dense types I and III collagen matrix. Capsular fibrosis and contracture have been suggested to stiffen the shoulder capsule and thus restrict range of motion^5^ similar to that in other fibrotic diseases such as Dupuytren’s disease. Fibroblasts are primarily responsible for the synthesis and remodelling of extracellular matrix and therefore tissue remodelling and repair^3^. Increasingly, activation or impaired fibroblast function has been associated with chronic inflammatory diseases such as RA^1^ and soft tissue diseases such as tendinopathy^1^. In normal circumstances fibroblasts act a sentinel like cells within tissue and are the first to respond to danger/damage signals^26^. Activated/damaged fibroblasts in healing tissues or inflamed can promote the recruitment and retention of immune cells as well as regulating their behaviour^12^. Pathogenic activated stromal fibroblasts have previously been identified in RA, these were characterised by increased expression of podoplanin (PDPN), CD106 (VCAM-1) and CD248 (tumour endothelial marker-1/endosialin) ^2^. PDPN is a transmembrane glycoprotein implicated in the invasiveness of cancer metastasis^34^ and VCAM-1 functions as a cell adhesion molecule^21^. CD248 is a transmembrane receptor with ligands that include collagen 1 ^22^ and is up-regulated by inflammation and fibrosis while CD146 (M-CAM) is a cell adhesion molecule that has been demonstrated to appear on a small subset of T and B lymphocytes in the peripheral blood of healthy individuals^9^. These stromal fibroblast activation markers have been identified in different locations of the synovium in rheumatoid arthritis^1^ and soft tissue tendinopathy^8^. In the tendon, inflammation induced stromal fibroblast activation which was more profound in diseased compared to healthy tendon cells. The authors suggest persistent stromal fibroblast activation is an important mechanism for the development of chronic inflammation in soft tissue musculoskeletal conditions drawing parallels with frozen shoulder.

We therefore hypothesised that subpopulations within the stromal compartment might become activated and expanded during the inflammatory processes occurring in frozen shoulder. We designed a study to investigate whether stromal activation markers might contribute to the pathogenic mechanisms of frozen shoulder and co-exist with inflammatory cytokines. The aims of this study were to evaluate stromal activation in patients presenting with frozen shoulder in an attempt to understand their association with this disease.

## Materials and Methods

### Study design

A prospective case-control study was conducted at the Institute of Infection, Immunity and Inflammation at the University of Glasgow, Scotland, UK. The study was conducted in accordance with ethics approval from Human Research Ethics Committee –West of Scotland REC (REC14/WS/1035).

### Inclusion and exclusion criteria

The study included patients with primary frozen shoulders who were receiving arthroscopic capsular releases. It also included patients with unstable shoulders undergoing arthroscopic stabilization as control groups as these patients were thought not to have their primary shoulder disease affecting the shoulder capsule and required glenohumeral arthroscopy, which was necessary to obtain tissue samples.

The diagnosis of frozen shoulder was made using Codman’s modified criteria by Zuckerman and Rokito^35^. The inclusion criteria for the frozen shoulder population comprised a painful stiff shoulder with insidious onset, restricted elevation to ≤100°, passive external rotation ≤50% of the normal contralateral shoulder, loss of function of the affected arm, pain at night and inability to lie on the affected side, and unremarkable radiographic and ultrasound findings of the glenohumeral joint. The exclusion criteria included history of previous surgery of the involved shoulder and radiologic or arthroscopic signs of fracture, glenohumeral arthritis, history of shoulder trauma and concomitant rotator cuff tear.

### Sample Collection

Tissue specimens were excised from the shoulder capsule by four different shoulder surgeons (NLM,DM,AA, UGF) during glenohumeral joint arthroscopy in patients undergoing either arthroscopic capsular release, stabilization or subacromial decompression surgery as previously described^19^. The tissue was removed by meniscal basket forceps from the shoulder capsule of the rotator interval. The biopsied tissue was immediately fixed in 10% buffered formalin for 18 to 24 hours, dehydrated, and embedded in paraffin wax.

### Immunohistochemistry

Samples for immunohistochemistry were immediately fixed in 10% formalin solution and stained with haematoxylin and eosin. Ten unstained sections were immunostained for the stromal activation markers CD248, CD146, CD90, PDPN, CD34, and FAP (all Sigma-Aldrich). The unstained sections were de-waxed in an oven at 60°C for 35 minutes. They were then immersed in 2 changes of xylene for 5 minutes each, and deparaffinated in 100%, 90% and 70% ethanol solution for 3 minutes of 2 cycles, thereafter they were rehydrated in distilled water for 5 minutes. Sections were then immersed in 0.5% hydrogen peroxide made up in methanol for 30 minutes to block endogenous peroxidase activity and then Tris Buffered Saline 0.05% Tween (TBST) solution (pH7.6). Endogenous peroxidase activity was quenched with 3% (v/v) H_2_O_2_, and nonspecific antibody binding blocked with 2.5% horse serum in TBST buffer for 30 minutes. Antigen retrieval was performed in 0.01M citrate buffer for 20 minutes in a microwave. The sections were left overnight to incubate at 4°C.

The sections were brought to room temperature for 30 minutes, and washed in TBST solution for 5 minutes. The sections were then incubated in DAB solution (1 drop DAB to 1ml DAB diluent) for 30 seconds, and quickly washed with TBST solution. They were then dipped twice in haematoxylin, washed in water, dipped in Scott solution to blue and then dehydrated in 70% ethanol, 90% ethanol for 30 seconds,1 minute respectively, and 2 changes of 3 minutes in 100% ethanol. Then they were cleared in 2 changes of xylene for 3 minutes and mounted in DPX with a coverslip.

### Cell culture and treatments

Control and diseased fibroblasts were explanted form capsule biopsies undergoing surgery as described above. Patients ranged from 18-30 (Table 1) undergoing elective arthroscopic stabilisation surgery. Cultures were maintained in a humidified environment at 37°C, 5% CO^2^ in RPMI 1640 supplemented with 10% heat-inactivated Fetal Bovine Serum (FBS), 100U/ml penicillin, 100µg/ml streptomycin and 2nM L-glutamine (all Thermo Fisher Scientific). Cells were grown to subconfluency and passaged using trypsin-EDTA (Sigma-Aldrich). Fibroblasts from the second and third passage were seeded two days prior to stimulation.

**Table 1.**

| Patient Demographics |  |  |
| --- | --- | --- |
|  | Adhesive Capsulitis | Control Capsule |
| No. of Patients | 10 | 10 |
| Mean Age (years), (range) | 52 (40-63) | 24 (18-30) |
| Gender (F:M) | 7:3 | 3:7 |
| Duration of symptoms (months) | 6.3 ± 3 | 3.9 ± 2 |
| Range of motion (degrees) |  |  |
| Forward flexion | 80 ± 12 | 160 ± 8 |
| Abduction | 62 ± 10 | 145 ± 10 |
| External Rotation | 11 ± 5 | 60 ± 15 |
| Mean number of steroid injections prior to surgery | 2.2± 0.2 | 0.8± 0.1 |

Fibroblasts were seeded at 2.5 x10^4^/ml in 24 well culture plates and allowed to rest for 48 hours. Fibroblasts were then stimulated with 10ng/ml recombinant human IL-1β (Biolegend) for 24 hours in complete RPMI.

### Microscopic Analysis

Tissue sections were examined under light microscopy at x10 and x40 magnifications and compared to the negative controls. Representative photomicrographs of the sections in each group were made. Two experienced examiners (JHR, NLM) blinded to the identity of the stained sections, performed the counting of each section.

Tissue analysis occurred in two stages: the first stage had all samples being given a semi-quantitative grade based on the percentage of positively-stained cells (taken over the total number of cells in that field) in 10 random high-powered fields. The following semi-quantitative grading was used: Grade 0, no staining, Grade 1, mild, ≤10% of cells stained positive; Grade 2, moderate, 10% to 20% of cells stained positive, Grade 3, strong, ≥20% of cells stained positive. The mean of these values was analysed by an unpaired Student’s *t*-test.

In the second stage, the samples had 10 random high-powered fields analysed at x40 magnification, and cells in each field were counted manually. The mean percentage of positively stained cells was taken over the total number of cells per high powered field, similarly, the results were analysed by an unpaired Student’s t-test.

### RNA extraction, cDNA synthesis and real time qPCR

Cells were placed in lysis buffer containing 1% 2-mercaptoethanol and RNA was extracted using mini columns according to the manufacturer’s instructions. RNA concentration and purity was determined using a spectrophotometer (Nanodrop 2000, Invitrogen). 100ng of RNA was converted to cDNA using High Capacity cDNA reverse transcription kit (Invitrogen) according to manufacturer’s instructions. cDNA was diluted 1 in 5 using RNase free water. qPCR was performed using PowerUp Sybr Green Mastermix (Invitrogen) and 1μl cDNA was used per reaction with 0.1 μM of forward and reverse primers. Each sample was run in duplicate and normalized to GAPDH or 18S endogenous control. Data represents fold change from untreated cells (2^-ΔΔCT^).

Primers (Integrated DNA Technologies) were as follows:

***GAPDH*** (f) 5′-TCGACAGTCAGCCGCATCTTCTTT-3′ (r) 5′-ACCAAATCCGTTGACTCCGA CCTT-3′
***CD248*** (f) 5′- CCCAAATCCCAAGGGAAGAT-3′ (r) 5′- CTGTGCTCGGCAAGACC-3′
***MCAM*** (f) 5′- CGGCACGGCAAGTGAAC-3′ (r) 5′- GCATTCAACACCTGTCTCCAAC-3′
***PDPN*** (f) 5′- CTTGACAACTCTGGTGGCA-3′ (r) 5′- GCGCTTGGACTTTGTTCTTG-3′
***VCAM*** (f) 5′- GCAAGTCTACATATCACCCAAGA-3′ (r) 5′- TAGACCCTCGCTGGAACA-3′
***IL-6*** (f) 5′- CAC TCA CCT CAG AAC GAA -3′ (r) 5′- GCT GCT TTC ACA CAT GTT ACT -3′
***IL-8*** (f) 5′- GTG CAT AAA GAC ATA CDC CAA ACC -3′ (r) 5′- GCT TTA CAA TAA TTT CTG TGT TGG -3′
***CCL20*** (f) 5′- GTC TTG GAT ACA CAG ACC GTA TT -3′ (r) 5′- GTG TGA AAG ATG ATA GCA TTG ATG T -3′

### Measurement of cytokine release by ELISA

Cell culture supernatants were collected and the concentrations of IL-6, IL-8 (both Invitrogen) and CCL20 (Biolegend) were determined using commercially available ELISA kits. Optical density was measured at 450 nm by a microplate reader.

### Statistical Analysis

Results are reported as the mean value and the standard error of mean. Comparisons between groups were made with two-way paired Student *t*-tests, Mann-Whitney U tests, and Kruskal-Wallis one-way analysis of variance on ranks. Based on our previous immunohistochemical studies^25^ and power calculations^6, 28^ (power of 0.8 and beta error of 0.2), we identified that each group required ten patient samples to detect a 20% difference in immunostaining for the various alarmin markers.

## Results

### Demographics

Shoulder capsule of a total of 20 patients were included in the study. Of these 20 patients, 10 had adhesive capsulitis (7 female, 3 male) with a mean age of 52 (range, 40-63 years) and 10 control subjects (shoulder instability, supraspinatus tendinopathy) (3 female, 7 male) with a mean age of 31 (range, 16-46 years). The control group was significantly younger than adhesive capsulitis group (p<0.01). The mean number of glenohumeral joint injections was significantly (p<0.05) greater in the frozen shoulder group compared to control (Fig. 1).

**Figure 1:**
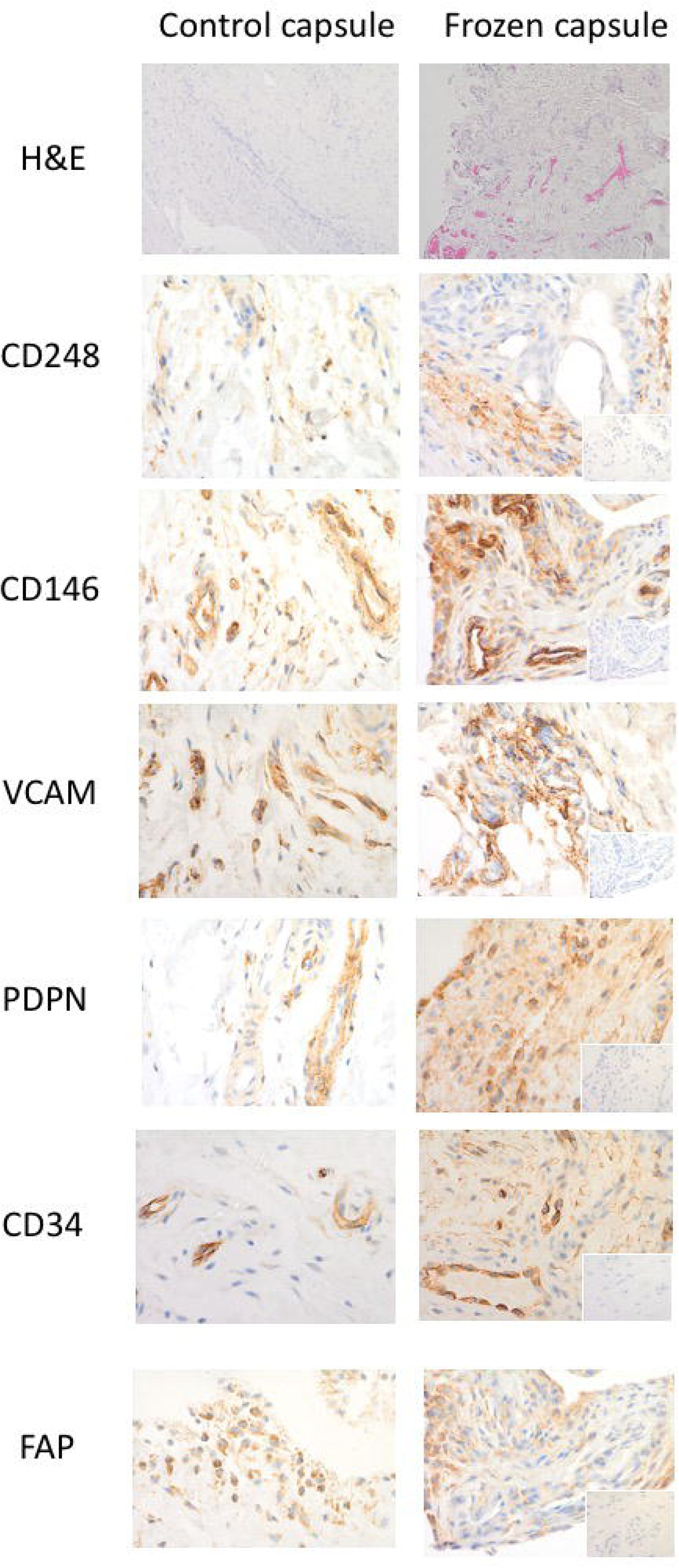
Stromal activation markers in shoulder capsule. Representative images of and control capsule and frozen shoulder tissue sections stained with Haematoxylin and Eosin (x10) and antibodies against CD248, CD146, VCAM, PDPN, CD34, and FAP (x40). Isotype control in bottom right corner (x10).

#### Pain and stiffness scores

Patient-ranked pain frequency (p=0.01), pain severity (p=0.03) and stiffness (p=0.03) were significantly higher in the frozen shoulder group compared to the instability group (Fig. 1). Shoulder motion was also significantly more restricted in external rotation, internal rotation, forward flexion and abduction in frozen shoulder patients compared to control group (Fig.1).

### Histologic appearance of the shoulder capsule

The histologic appearance of the H&E-stained frozen shoulder capsule specimens showed densely packed collagen fibres and fibroblastic proliferation within the fibrous stroma compared with the control (Fig 2). There were large numbers of capillaries and venules in the subsynovium of frozen shoulder samples compared with the controls. Capsule specimens from 5 control patients (3 instability, 2 supraspinatus tendinopathy) also exhibited few capillaries and venules in the subsynovium whereas 4 (3 instability, 1 supraspinatus tendinopathy) had increased vascularity.

**Figure 2:**
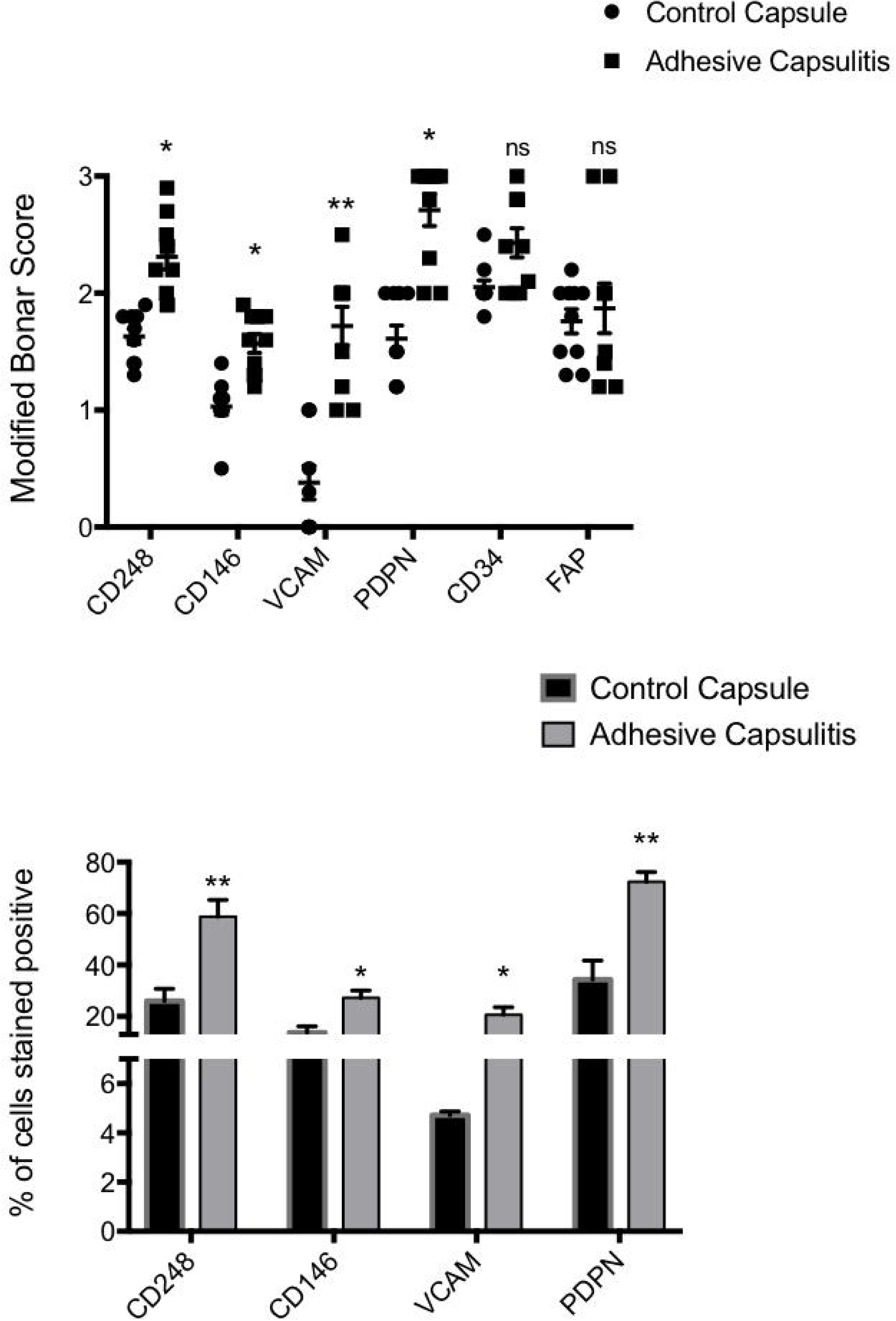
Stromal activation marker expression in control capsule and frozen shoulder. Graphs illustrate Modified Bonar scoring & % of cells positively stained for samples of human capsule biopsies for expression of CD248, CD146, VCAM, PDPN, CD34 and FAP, mean ± SEM, n = 10 for control capsule, n = 10 for frozen shoulder capsule. Modified Bonar scoring system depicts mean score per sample based on 10 high power fields. 0 = no staining, 1 = <10%, 2 = 10–20%, 3 = >20% + ve staining of cells per high power field. *p < 0.05, **p < 0.01 (Students *t-*test).

### Diseased capsule tissues express stromal fibroblast activation markers

There was significantly greater expression of the stromal activation markers, CD248, CD146, VCAM and PDPN in the frozen shoulder group compared with the control group (p < 0.05) (Figs. 1&2). CD248, CD146 and VCAM were mainly localised to stromal cells (Fig. 1) however, whilst exhibiting a similar stromal pattern, PDPN also showed significant extracellular matrix staining pattern. Further sub analysis of tissue distribution revealed that PDPN was most abundantly expressed in adhesive capsulitis with approximately 60% of cells positive versus 40% in control tissues (Fig. 2) followed by CD248 (60% versus 25% control, p<0.01), VCAM (20% versus 5% control, p<0.01) and CD146 (25% versus 18% control, p<0.05). We found no differential expression of CD34 nor FAP between normal of diseased tissue samples.

### Inflammatory cytokines are overexpressed in diseased versus normal capsular cells

To understand if the presence of stromal activation markers was associated with an inflammatory signature within frozen shoulder fibroblasts we analysed the gene expression of inflammatory associated cytokines/chemokines. We found that fibroblasts cultured from diseased capsule produced greater levels of inflammatory proteins compared to cells cultured from control capsule (Fig. 3). This was the case for the cytokines IL-6 and IL-8, as well as the chemokine CCL20.

**Figure 3:**
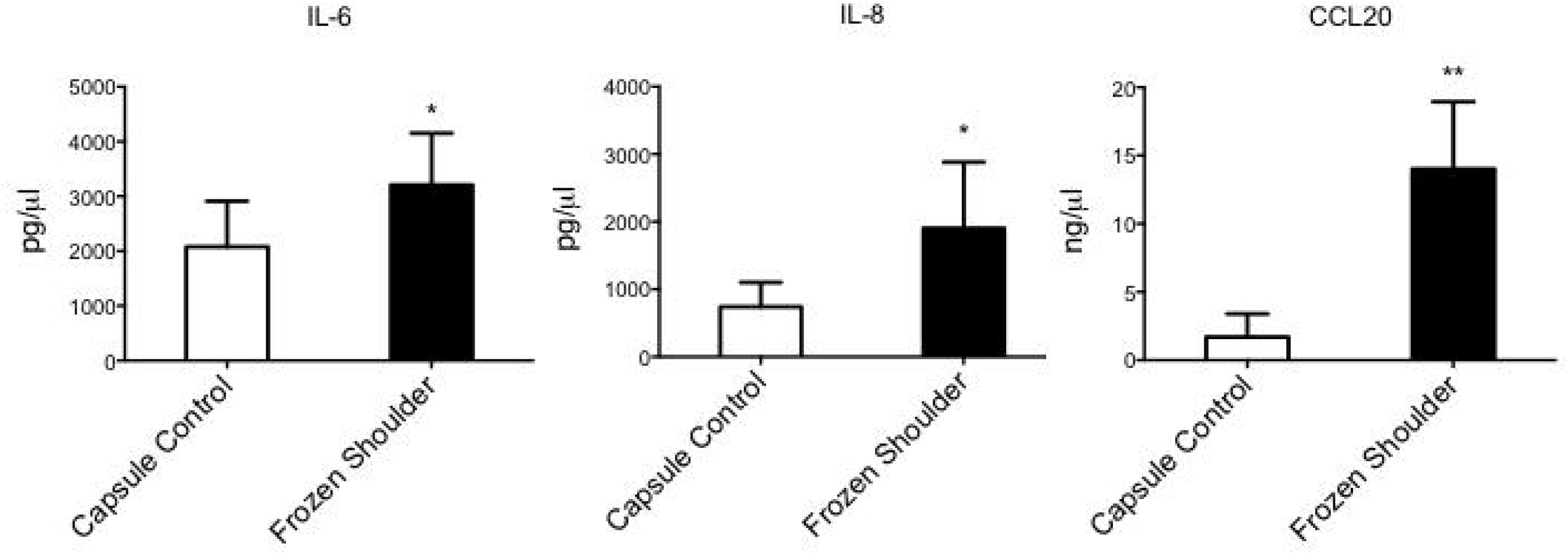
Inflammatory cytokine production from capsule derived fibroblasts. IL-6, CCL20 and IL-8 secretion from healthy capsule and diseased capsule derived fibroblasts, mean ± SEM, n=5 *p < 0.05 (Students *t*-test).

### Inflammation induces persistent stromal activation in capsule-derived stromal cells

Having identified that stromal fibroblast activation markers were expressed at higher level in diseased capsular tissues and stromal cells cultured from these tissues secreted more inflammatory proteins we sought to investigate whether an inflammatory stimuli (IL-1β) could induce stromal fibroblast “memory” in healthy cells *in vitro*. Figure 4 demonstrates that healthy fibroblasts had increased CD248, PDPN and VCAM mRNA expression compared to unstimulated cells (*p*<0.05). Furthermore, IL-1β treatment for 24 hours also induced CD248, PDPN VCAM *and* CD146 protein expression in healthy capsule cells (Fig 4). To ascertain whether this inflammatory simulated “diseased phenotype” fibroblast had any functional relevance we measure the inflammatory proteins released into the supernatant. We found control cells following inflammatory stimuli significantly increased mRNA expression and protein secretion of IL-6, IL-8 and CCL20 (Fig 5).

**Figure 4:**
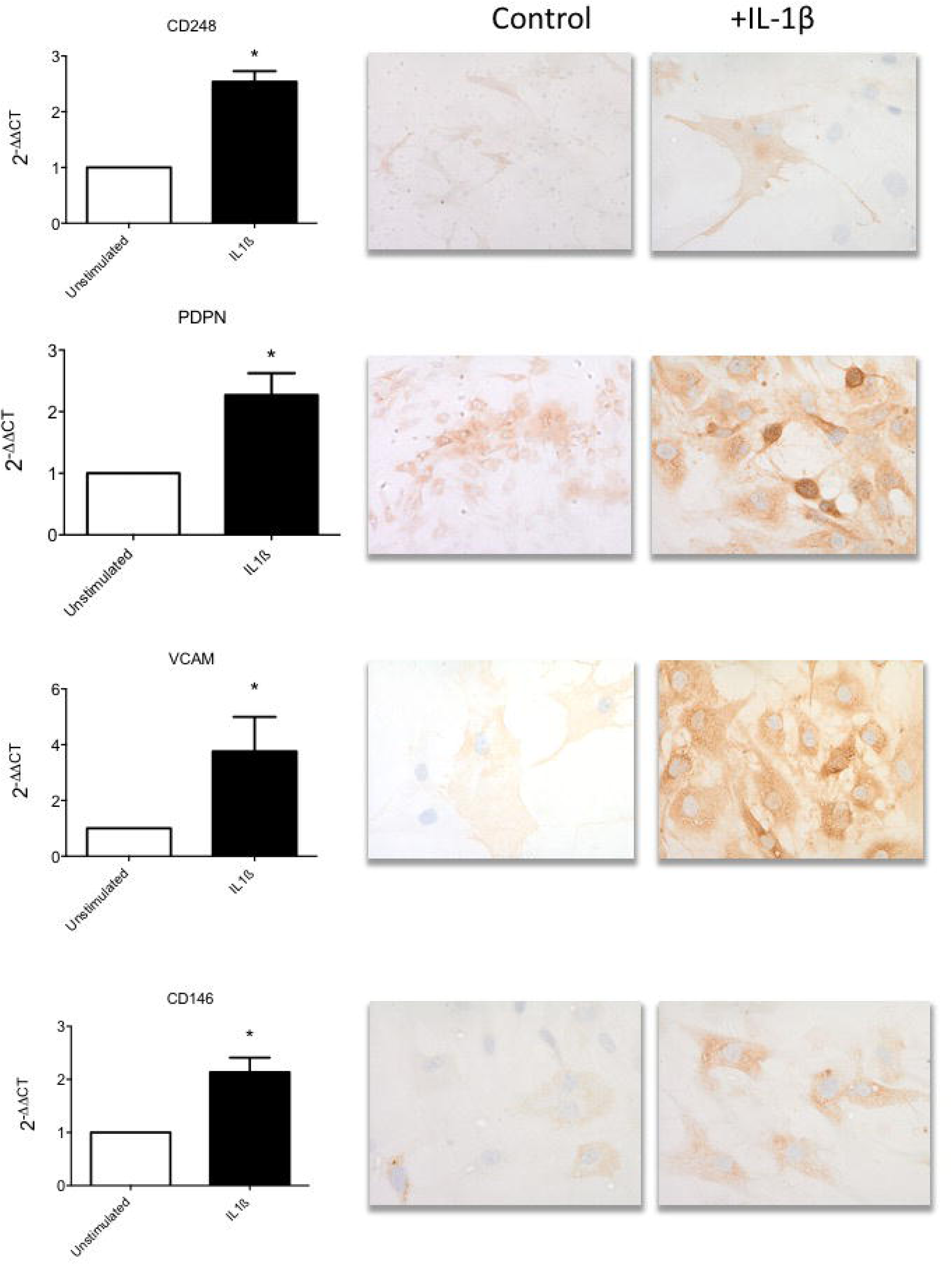
Stromal activation markers are upregulated in shoulder capsule upon inflammatory stimulus. and inflammatory protein in healthy fibroblasts following IL-1β (10ng/ml) stimulation. (a) Stromal activation marker (CD248, PDPN, VCAM & CD146) mRNA expression following IL-1β stimulation of cultured healthy derived fibroblasts. Data represented 2^-ΔΔCT^, normalized to GAPDH housekeeping control, mean ± SEM, n=5. *p < 0.05, **p < 0.01 compared to control samples (Students *t*-test). (b) Representative images of healthy capsule derived fibroblasts following IL-1β stimulation. Cells stained with antibodies against CD248, PDPN, VCAM and CD146, image at x40.

**Figure 5:**
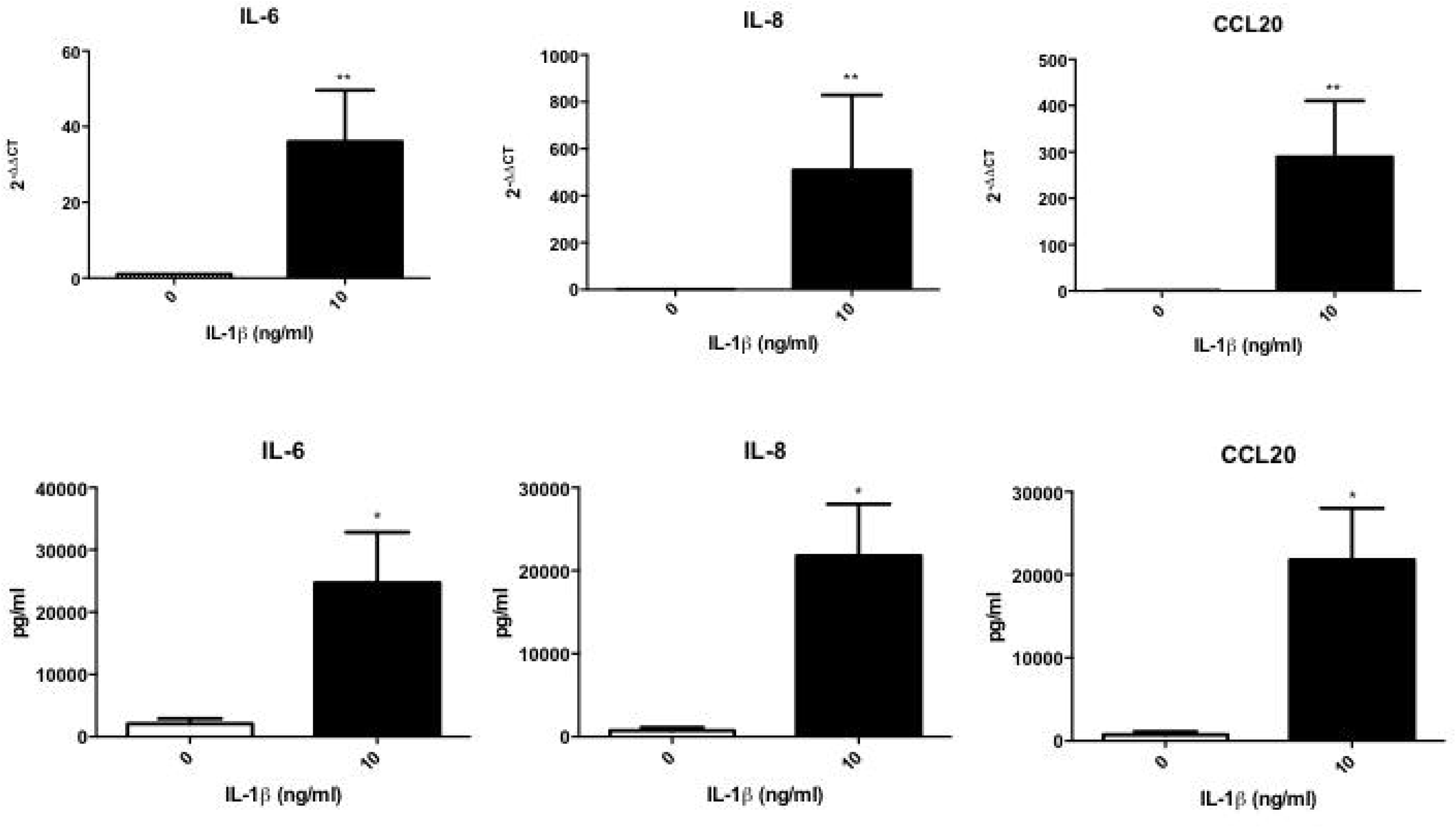
Activated capsule fibroblasts drive proinflammatory cytokine production. IL-6, CCL20 and IL-8 mRNA and protein expression following IL-1β stimulation. mean ± SEM, n=5, *p < 0.05, **p < 0.01 compared to control samples (Students *t*-test).

**Figure 6:**
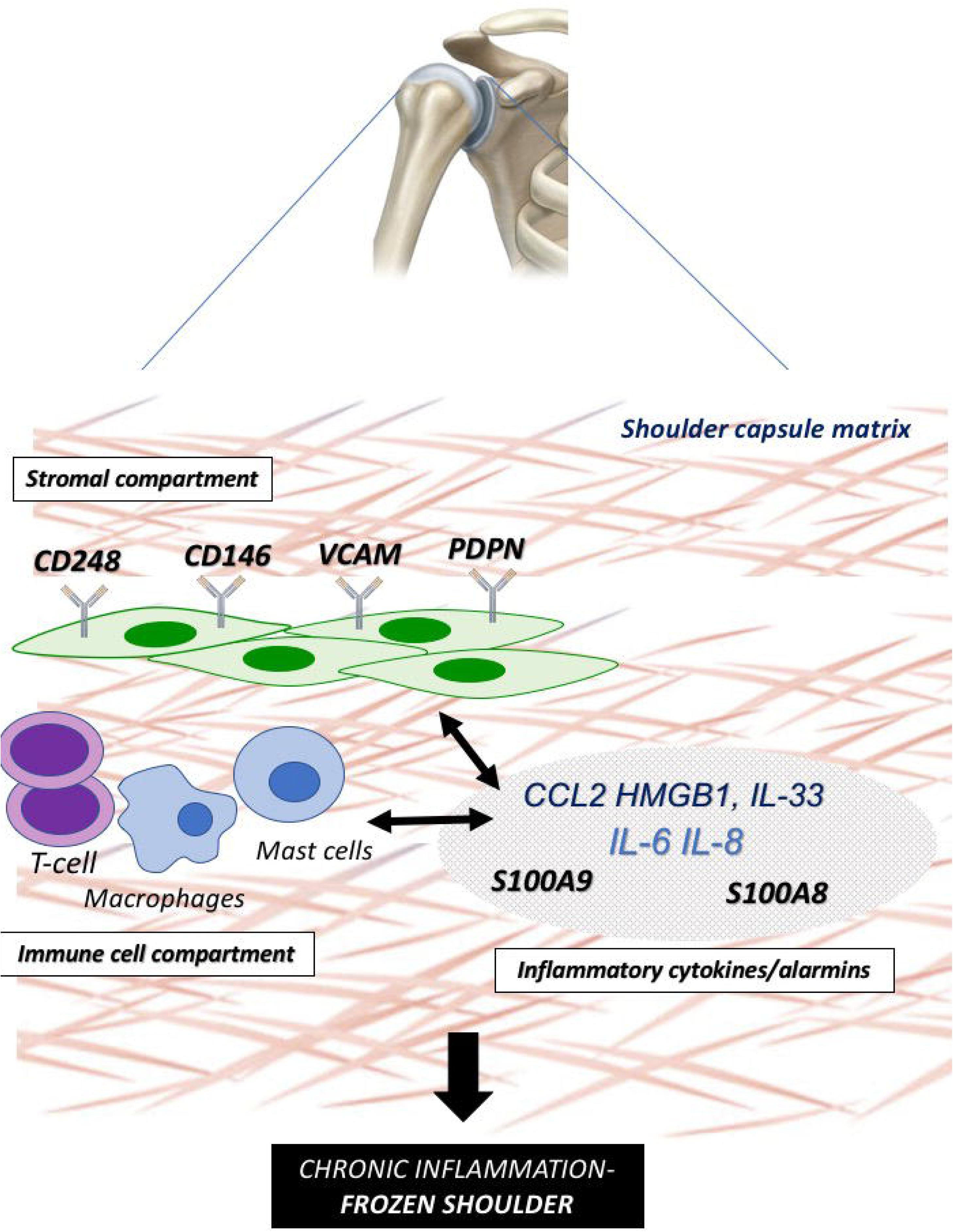
Immunobiology in frozen shoulder. Schematic illustration representing the functional interplay between the “activated” stromal fibroblast (CD248+,CD146+,VCAM+,PDPN+) (stromal compartment), infiltrating and resident immune cells (immune compartment) and downstream cytokine/alarmin dysregulation interacting to drive a proinflammatory phenotype with aberrant tissue fibrosis in the frozen shoulder joint capsule.

## Discussion

This study is the first to investigate the expression of stromal activation molecules in frozen shoulder capsule and has demonstrated the presence of CD248, CD146, VCAM, PDPN, CD34 and FAP in shoulder capsule tissue of patients with frozen shoulder co existing with proinflammatory cytokine expression. Importantly inflammatory activation of healthy fibroblasts leads to overexpression of stromal activation markers in addition to secreting inflammatory proteins suggesting this may drive disease chronicity.

Increasingly datasets from human biopsy studies in patients with frozen shoulder have suggested a strong immune component to the underlying disease process^31^. Histologically, frozen shoulder tissue is characterized by fibroblasts, myofibroblasts and chronic inflammatory cells, including mast cell, T-cells, B-cells and macrophages^4, 17^. Furthermore cytokines such as IL-1β, IL-6 and TNF-α, which are known to drive inflammatory/matrix interactions^27^ including fibroblast activation and dysregulated collagen synthesis are upregulated in adhesive capsulitis versus normal shoulder capsule^30^ and the subacromial bursa^20^ of frozen shoulder patients. The most abundant cell type within tissue stroma remains the fibroblast which play a central role in determining the site at which inflammation occurs, and influence the persistence of the inflammatory process^33^. Once activated, fibroblasts have been shown to produce TNFα, IL-1, and IL-6, cyclooxygenase-2, the polysaccharide hyaluronan, as well as inflammatory chemokines^11^ thus sustaining leukocyte recruitment in to the inflamed synovium. Podoplanin (PDPN), a 38-kDa type I transmembrane glycoprotein which is found in some stromal fibroblasts, and an abundance of PDPN-positive stromal fibroblasts is associated with poor prognosis in lung adenocarcinoma, invasive breast cancer, and esophageal squamous cell carcinoma patients^34^. Furthermore, PDPN expression can be upregulated by a number of pro-inflammatory cytokines, including IL-22, IL-6, IFN-γ, TGF-β, IL-1β, and TNF-α, but the signalling pathways involved are largely unknown. Given the previous finding that PDPN may be a component of chronic soft tissue inflammation our results demonstrating significantly increased PDPN in frozen shoulder suggest that the chronic inflammation changes may be similar to those seen in tendinopathy. Further work is now required to dissect the role of podoplanin in immune cell interactions within frozen shoulder and its role in driving a dysregulated fibrotic tissue response.

VCAM-1 belongs to the Ig superfamily and is expressed on monocytes, endothelial cells and synovial cells. In inflammatory reactions VCAM-1 is a surface glycoproteins that promote adhesion and subsequent recruitment of leukocytes. The levels of circulating VCAM-1 in plasma and synovial fluid both were significantly increased in rheumatoid arthritis patients compared to normal controls^23^. VCAM-1 has also been shown to be upregulated following damage associated protein (alarmin) release from epithelial cells. In a similar cohort of patients we have recently reported elevation of the alarmins HMGB1 and IL-33 and S100A8/A9 in the shoulder capsule of patients with frozen shoulder compared to controls while furthermore confirming significantly increased neoinnervation linked to patient reported pain^7^. Thus, given that activation of the immune system after tissue injury is partially due to the interactions of alarmins released from necrotic cells and subsequent upregulation of stromal activation markers this association may represent an important synergy between stromal markers and inflammatory molecules that can drive the disease chronicity and fibrosis associated with frozen shoulder. Future work assessing associations between stromal markers and alarmin molecules in patient subgroups may help address whether these interactions are linked to failure of tissue resolution. Interestingly, agents targeting stromal biology have delivered promising *in vitro* and *in vivo* data^10, 29, 32^ particularly altering matrix regulation. This suggests targeting stromal activation may offer novel therapeutic potential in frozen shoulder.

In contrast to other stromal associated pathologies FAP and CD34 were expressed to the same extent in frozen shoulder when compared to control capsule. FAP was initially identified in the context of RA^13^ with activated phagocytes expressing this protein in the inflammatory lesions in synovium^14^. Thus, in frozen shoulder it appears that the FAP and CD34 may not represent activation markers but constituently expressed surface proteins which may be involved in homeostatic tissue responses.

There are limitations inherent in our study. The study had a small sample size however our previous studies^15, 19, 28^ and power analysis confirms that 10 patient samples per group gives us confidence that the changes seen do represent a biologically relevant change in protein expression. Additionally, there was selection bias in that we only studied patients with frozen shoulder that warranted a surgical procedure and thus did not include asymptomatic patients or patients with less pain intensity with frozen shoulder. Furthermore, the patients in the control group were younger, predominately male, and had shorter duration of symptoms compared with the frozen shoulder group which is an unfortunate consequence of the availability of human control tissues. Three surgeons were involved in the collection of samples, giving rise to the possibility that surgical technique and biopsies had some user-related differences. Despite this, the same method of obtaining samples was used in order to reduce the surgeon-related bias. This study ensured that comparison only occurred between shoulder capsule samples of patients with adhesive capsulitis and shoulder capsules samples of patients with instability. This allowed a comparison between the same type of tissue and removed the possibility that pathology in other tissue types – such as tendons, could confound the study. The exclusion of patients with concurrent pathologies such as osteoarthritis and rheumatoid arthritis also allowed better isolation of cases to adhesive capsulitis in the shoulder. As such, results of immunoreactivity were less likely to be affected by other comorbidities.

## Conclusion

This study has shown for the first time that there is elevation of stromal activation markers in the shoulder capsule of patients with frozen shoulder compared to control capsular tissue while furthermore confirming significantly increased proinflammatory signalling within frozen shoulder capsule. This data is consistent with the hypothesis that persistent stromal activation may play a role in the pathology of frozen shoulder and could explain the cellular mechanisms behind capsular fibrosis and persistent inflammation.

## Funding

This work was funded by the Medical Research Council (MR/K501335/1 and MR/R020515/1, Arthritis Research UK (21346) and Tenovus Scotland.

## Contributions

M.A, M.M and N.L.M. conceived and designed the experiments. M.A, M.M, E.G.M, P.M, , L.A.N.C, and J.H.R performed experiments. I.B.M,U.G.F, D.M and A.A, provided expert advice. All authors analysed the data. M.A and N.L.M wrote the paper.

## Competing Interests

The authors declare no competing interests.

## Ethical approval information

All procedures and protocols were approved by the Ethics Committee under approval numbers West of Scotland REC (REC14/WS/1035) with informed consent obtained and carried out in accordance with standard operative procedures.

## Data sharing statement

M.A and M.M. have access to all the data and data are available upon request.

